# Mistle: bringing spectral library predictions to metaproteomics with an efficient search index

**DOI:** 10.1101/2022.09.09.507252

**Authors:** Yannek Nowatzky, Philipp Benner, Knut Reinert, Thilo Muth

## Abstract

**Motivation:** Deep learning has moved to the forefront of tandem mass spectrometry-driven proteomics and authentic prediction for peptide fragmentation is more feasible than ever. Still, at this point spectral prediction is mainly used to validate database search results or used for confined search spaces. Fully predicted spectral libraries have not yet been efficiently adapted to large search space problems that often occur in metaproteomics or proteogenomics.

**Results:** In this study, we showcase a workflow that uses Prosit for spectral library predictions on two common metaproteomes and implement an indexing and search algorithm, Mistle, to efficiently identify experimental mass spectra within the library. Hence, the workflow emulates a classic protein sequence database search with protein digestion but builds a searchable index from spectral predictions as an in-between step. We compare Mistle to popular search engines, both on a spectral and database search level, and provide evidence that this approach is more accurate than a database search using MSFragger. Mistle outperforms other spectral library search engines in terms of run time and proves to be extremely memory efficient with an 8 to 22-fold decrease in RAM usage. This makes Mistle universally applicable to large search spaces, e.g. covering comprehensive sequence databases of diverse microbiomes.

**Availability:** Mistle is freely available on GitHub at https://github.com/BAMeScience/Mistle.

## 1 Introduction

Metaproteomics is a key technology for characterizing proteins in complex microbial samples at a given time point (Wilmes and Bond, 2004) and can provide pivotal information about taxon-specific functional activity, as well as signaling and metabolic pathways within the microbial community (Hettich *et al*., 2013; Tanca *et al*., 2017). This enables studying health and disease cases of host species, and ecological dynamics in all kinds of biological systems and microbiomes (Callieri *et al*., 2018; Hettich *et al*., 2013; Scholz *et al*., 2015). Inherent to the proteomic investigation of microbial communities is an enlarged search space, as many species, often previously unknown ones, are present in a typical microbiome sample and need to be queried (Schiebenhoefer *et al*., 2019).

Peptide identification lies at the heart of high-throughput proteomics workflows, where the collected sample of proteins is usually subjected to enzymatic digestion and liquid chromatography (LC) coupled with tandem mass spectrometry (MS/MS) (Coon *et al*., 2005; Hettich *et al*., 2013). Various search algorithms have been designed to identify the underlying peptides from the MS/MS spectra in a protein sequence database, assigning quality scores for so-called peptide spectrum matches (PSMs) (Verheggen *et al*., 2020). However, distinguishing true identifications from false positive hits becomes increasingly hard with a growing number of candidate peptides to match to, i.e. size of protein database that is used as a reference for peptide identification (Verbruggen *et al*., 2021). A greater number of potential matches make it statistically more likely that a random false match receives a higher score than the true match (Nesvizhskii, 2010; Verbruggen *et al*., 2021). When filtering for false positives, an increase in database size may lead to a reduced number of peptides identified. Even multi-stage search strategies, which aim to reduce the search space by tailoring the database through multiple search steps, may at the same time invoke more false discoveries (Muth *et al*., 2015; Verheggen *et al*., 2020). Evidently, there is a demand to overcome this inherent weakness of database search when facing large search spaces.

Due to the diversity of species and genera found in microbial communities, metaproteomic studies are especially resource-straining for common database search engines, such as MSFragger (Kong *et al*., 2017), which compare MS/MS scans to theoretical fragment ions of peptides in the sequence database. This is noticeable not just in terms of reduced sensitivity in peptide identification, but also in increased run time, and even more prominently in memory requirements by the algorithms, due to the large candidate spaces.

Machine learning approaches have been implemented to enhance peptide identification, e.g. by post-processing the database search results (Verbruggen *et al*., 2021). A particularly prominent example of this is Percolator (Käll *et al*., 2007), which uses semi-supervised learning with support vector machines. More recently, deep learning models such as Prosit (Gessulat *et al*., 2019) predict complete mass spectra including fragment intensities and retention time from peptide sequences, and thus offer a method to rescore database search results based on these predicted spectral features, coupled with Percolator.

Consequently, Prosit also provides the means to predict spectral libraries for entire proteomes, which can then be queried. As of now, metaproteomics has not yet had a chance to fully make use of such comprehensive prediction workflows covering the proteomes of many species, as this leads to massive spectral libraries. Current spectral library search software, such as SpectraST (Lam *et al*., 2007), is not equipped to meet run time and memory constraints imposed by such large MS/MS databases, covering *>*10,000,000 peptide spectrum predictions. At the same time, metaproteomics could benefit from more precise search algorithms, as the large search space has been shown to reduce sensitivity and exacerbates challenges with false discovery estimation in large metaproteomic settings (Muth *et al*., 2015). In fact, Gessulat *et al*. (2019) mention the use case of Prosit for metaproteomics, and manage to improve database search results by rescoring the top Andromeda hits (Cox *et al*., 2011) using the spectral predictions. In 2021, Verbruggen *et al*. presented a solution for large search spaces in proteogenomics, for ribosomal profiling, by using predicting spectral features to enhance identification rate and stringency in PSMs. However, to this day there is no efficient workflow to apply complete spectral library predictions to metaproteomics and efficiently search such substantial amounts of MS/MS data, to the best of our knowledge.

We propose such a workflow using an entire predicted library and directly search for the best matching peptide using spectral similarity measures. First, we digest the complete metaproteome sequence database (*in silico*) with EncyclopeDIA (Searle *et al*., 2020) and then use Prosit to predict MS/MS spectra for every peptide and charge configuration that is reasonably likely to occur in an MS/MS run. Finally, we use our novel **M**etaproteomic **i**ndex and **s**pec**t**ral **l**ibrary search **e**ngine, short Mistle, to query the spectral library.

Mistle creates a small, searchable index and is extremely run time and memory efficient. We achieve this by adapting the fragment index of MSFragger to spectral intensity matching. Additionally, we introduce an advanced index partitioning and query scheduling method to the algorithm and add hardware optimization, such as SIMD intrinsics in combination with multithreading, to greatly reduce memory footprint and run time.

This workflow virtually turns the sequence database search problem into a spectral library search problem. We benchmark the algorithmic performance of Mistle with state-of-the-art methods and examine the potential of our workflow to qualitatively and quantitatively improve metaproteomic studies on the peptide identification level, based on two sample metaproteomes, the lab-assembled nine-organism microbial mixture (9MM) by Tanca *et al*. (2013) and the extended simplified human intestinal microbiota (SIHUMIx) sample by Krause *et al*. (2020).

## 2 Methods

Mistle is inspired by the fragment index data structure introduced by Kong *et al*. in 2017 and employed in MSFragger. Instead of iteratively matching experimental spectra with every theoretical spectrum calculated from candidate peptide sequences, MSFragger constructs an index that stores theoretical fragment ions in an easily searchable format, enabling fast and simultaneous peak matching for peptide candidates. We adapt the core idea to the spectral search problem, where instead of a protein sequence database a predicted MS/MS library is queried.

While the main idea remains the same, i.e. searching fragments in the fragment index and updating scores of their parents, which are now MS/MS spectra rather than peptides, there are significant additional challenges to overcome. For one, peak intensities must be considered and stored in the fragment index. This immediately makes the fragment index larger, posing tangible problems for metaproteomics libraries, and slower to process as intensities need to be multiplied. Simply counting and summing up intensities, as it is the case for MSFragger, is no longer sufficient. Also, the index needs to be constructed from a spectral library, which is data-intensive to a point where it is infeasible to hold all data in random access memory (RAM). Thus, information required to construct the data structures needs to be carefully and continuously conveyed throughout the reading process to produce the final index.

Here, we introduce algorithmic solutions facing all these hurdles and propose optimizations to counteract increased run time arising from the additional multiplication-operations when matching peaks. Mistle is implemented from scratch in C++ 20 and features single instruction multiple data (SIMD) extensions.

### 2.1 Data structures

#### 2.1.1 Precursor index

Similarly to MSFragger, we require an auxiliary data structure, referencing all library targets, i.e. peptides linked with their predicted MS/MS spectra. We call it precursor index, as entries are searchable by the precursor peak. It is the equivalent to the peptide index described by Kong *et al*. (2017).

Specifically, the precursor index stores a unique identifier (ID, 32-bit unsigned integer) for every mass spectrum in the library, which each corresponds to exactly one peptide. The IDs are ordered by the precursor’s charge and mass-to-charge ratio (m/z). This serves as a reference for the fragment index. Additionally, a mapping must be provided from the ID to the rank of the spectrum in the precursor index. This is the inverse of the sorted precursor IDs. We implement this as an additional lookup vector, precomputed at index construction by a linear scan over the ranked ID vector.

#### 2.1.2 Fragment index

In the fragment index, all fragment ions *f*, i.e. peaks, of library MS/MS spectra are stored in form of triplets, of the ion mass (m/z value, *mz*_*f*_), peak intensity (*I*_*f*_) and the unique ID of the underlying parent spectrum: *f* = (*mz*_*f*_, *I*_*f*_, ID_parent_(*f*)). The latter provides a reference from fragment to the matched peptide/spectrum and facilitates an efficient search of peaks of matching candidate spectra.

For that to be possible, the fragments (triplets) are placed into bins based on their ion mass, given an adjustable bin width *B*. Inside each bin, fragments are sorted according to the parent rank in the precursor index, accessed via the parent ID. This way, for a certain mass range [*M, M* + *B*), every peak with *mz*_*f*_ ≥ *M* and *mz*_*f*_ *< M* + *B* of the entire spectral library is stored in the corresponding fragment bin. The order of those peaks allows for a division based on parent mass. The structure of the precursor and fragment index are illustrated in Figure 1.

**Figure 1:**
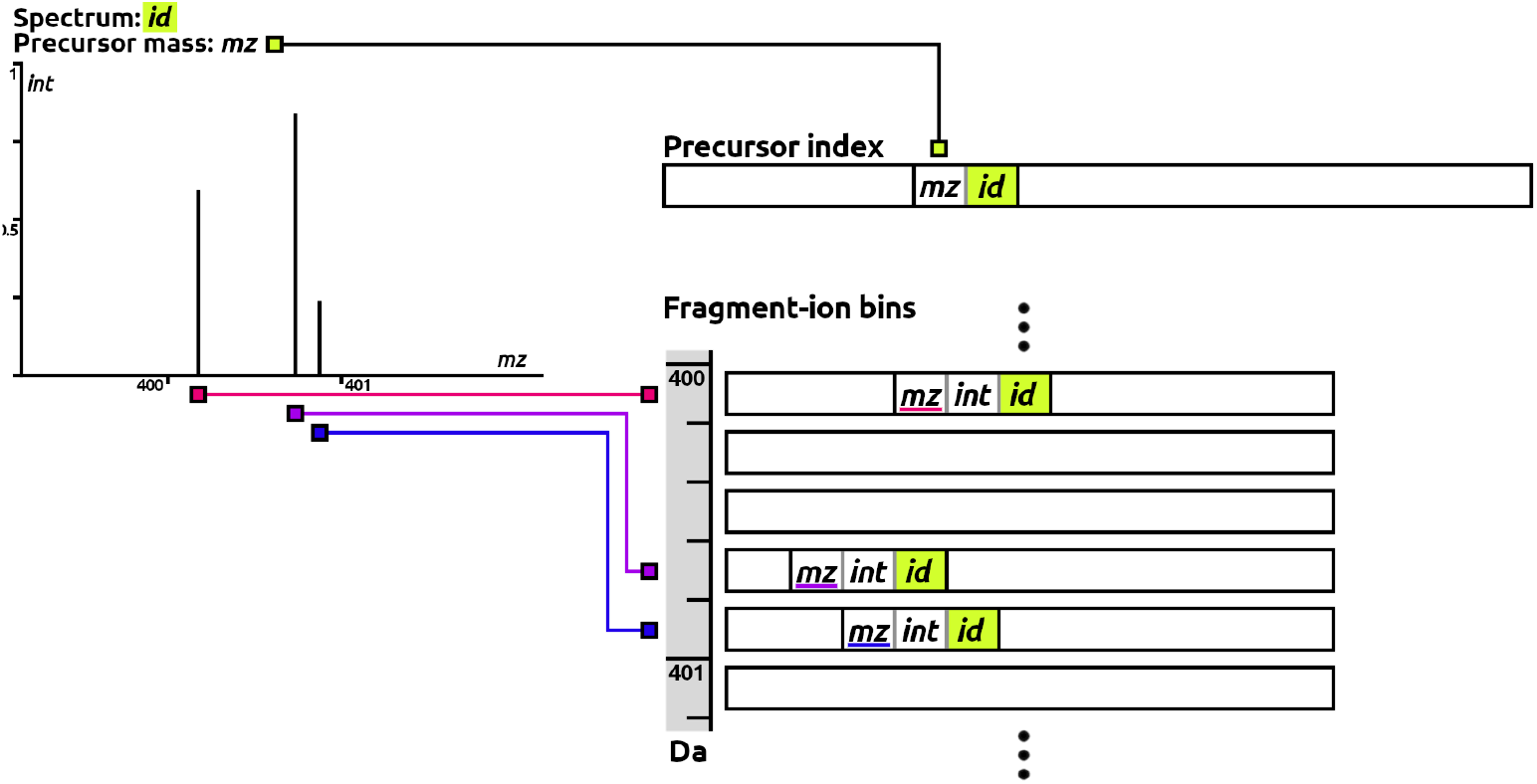
Precursor and fragment index data structures at construction. For an exemplary library spectrum (top left), it is shown how the precursor is included in precursor index sorted by mass (*mz*) and referenced by a unique identifier (*id*). All peaks are integrated in the fragment-ion bins corresponding to their ion mass and encoded as triplets (m/z value (*mz*), intensity (*int*) and parent ID (*id*)). Their order inside each bin is determined by the rank of the parent in the precursor index.

#### 2.1.3 Partitioning

As the union of precursor index and fragment index holds about as much information as the entire spectral library, the index space required grows linearly with the database size and needs to fit into main memory for efficient access. To make a search feasible for large reference libraries, we propose partitioning the main part of the index, i.e. fragment index, into several smaller sub-indices or partitions. Such a technique has been shown to be quite effective for other bioinformatic problems, showcased for instance by the DREAM index framework (Dadi *et al*., 2018). Ideally, each query spectrum only needs to be searched in one or a small number of partitions, which combined retain the original index data structure.

We achieve this by creating separate fragment bins for each partition, which we tie to non-overlapping precursor m/z intervals. A fragment triplet is stored in its corresponding fragment bin, for the partition only, where the parent’s precursor peak falls into the m/z interval. Each partition has the full number of fragment bins, and acts as an individual fragment index. This way, a query spectrum only needs to be searched in a partition matching its precursor mass, within a given m/z tolerance. Also, within each partition the search algorithm can be performed identically. Merely the number of library spectra included is reduced for each partition. This not only reduces physical space that needs to fit into the main memory at a time, but also the search space for a given query within the partition. Fewer comparisons are needed during the binary search, explained in section 2.2.

Note that this partitioning method is different from splitting the reference database, as offered by FragPipe for MSFragger. Instead of creating multiple indices for parts of the database, each partition represents a fraction of the main index and needs to be queried only once.

#### 2.1.4 Continuous index construction algorithm

As mentioned before, the input library might be arbitrarily large and in no particular order. When reading the data, the precursor index, which is necessary to order all fragments, is unknown, up until the very end. A practical, memory efficient approach is to create preliminary (unsorted) index partitions on the disk when reading the library and update the partitions once all relevant information has been obtained. A detailed description of the process can be found in the supplementary text.

### 2.2 Search Algorithm

Partitioning the fragment index creates an initial overhead when searching experimental spectra, as spectral queries need to be scheduled to relevant partitions and merged afterwards. This is performed by assigning each experimental query spectrum a unique identifier and constructing a list of query IDs for each partition to address, based on the precursor m/z and mass tolerance. Then, each partition with at least one query scheduled is loaded into main memory, and spectral matching is performed.

Initially, matches are ranked by the spectral dot product of normalized intensities, as described in Eq. (2) by Lam *et al*. (2007). The similarity scoring function is refined later on, described in detail in the supplementary text. Raw intensity values are square rooted before normalization to de-emphasive dominant peaks. By definition, a peak only contributes to the dot product, if a matching peak from the other spectrum exists in the same m/z bin. Hence, dot products to library spectra that are altered by matching that peak, must have a fragment entry in the corresponding fragment bin. We leverage the data structures from section 2.1 to perform a fast search, reminiscent of MSFragger’s fragment index search, but compute the spectral dot product in the process, as is illustrated in Figure 2.

**Figure 2:**
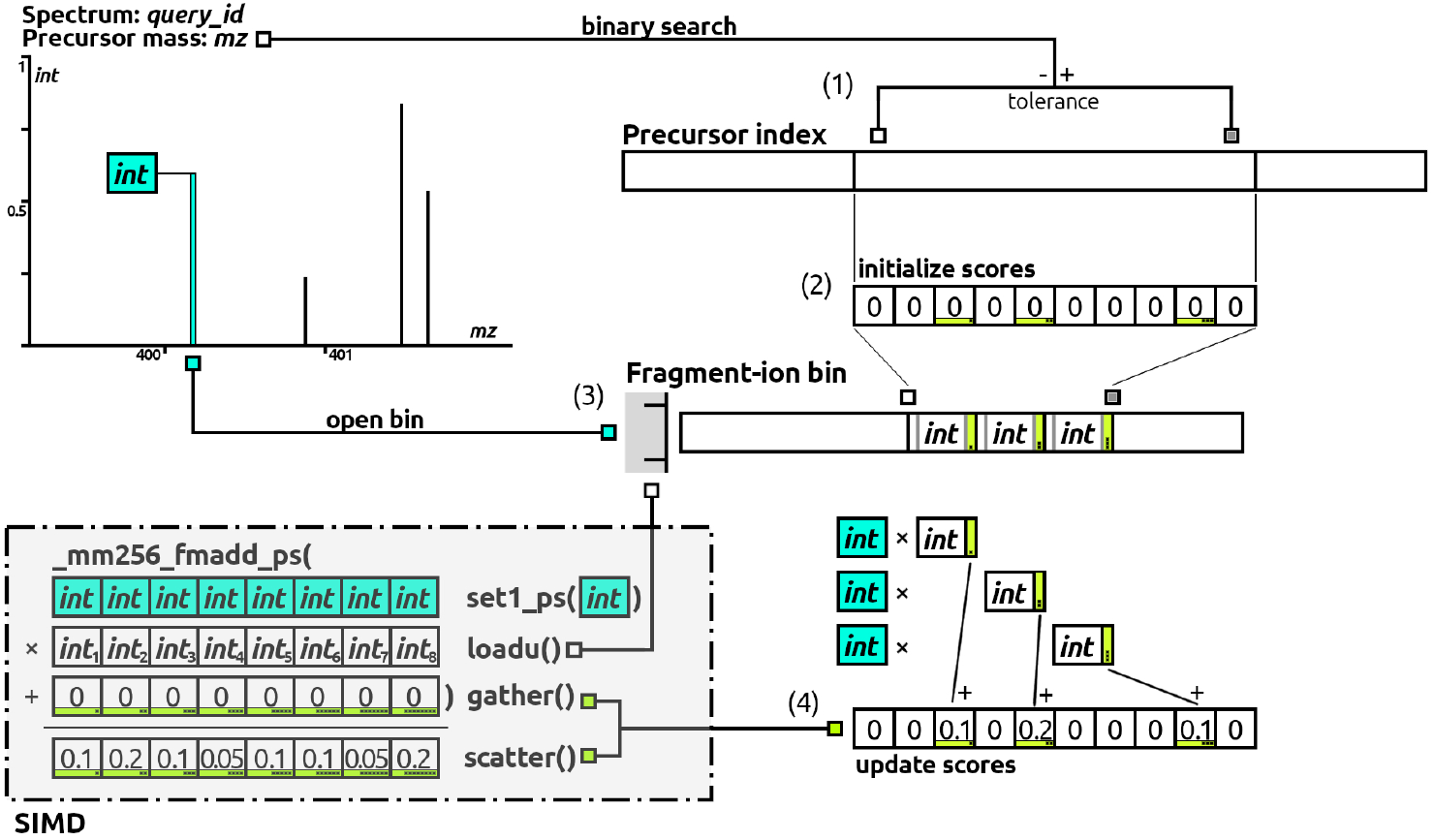
Illustration of the search process: matching an exemplary query spectrum (top left) to all indexed library spectra. First, the binary search step (1) is shown on the precursor index, where the lower and upper bound of candidate spectra is determined and an empty scoring vector is initialized (2). Thereafter, the peak-by-peak matching is shown for the fragment index, highlighted for the first query peak (turquoise) and the corresponding fragment bin. (3) Here, a binary search is performed to determine relevant matches with a parent rank within the lower and upper bound. (4) Lastly, fragment intensity entries are multiplied to the peak intensity and added to the respective parent scores. SIMD intrinsics may replace step 4, as shown on the bottom left, e.g. computing 8 intensity products and adding them to the dot products in a single CPU operation.

The overt novelty lies within matching fragments by their intensities, in addition to the m/z dimension, when iterating a fragment bin. The intensity product (*I*_*p*_*I*_*f*_) of query peak *p* = (*mz*_*p*_, *I*_*p*_) and fragment *f* is added to the parent score, which is accessed by the parent identifier ID_parent_(*f*). As this is computationally costly, we speed up the arithmetic operations using SIMD extensions. The fused multiply-add operation (in C++: mm256 fmadd ps), available for the Advanced Vector Extensions AVX2 and AVX512 architectures, allows parallel multiplication and addition of 8 to 16 floating-point numbers in a single CPU instruction. A schematic workflow with the use of SIMD for a 256-bit register is depicted in Figure 2, bottom left. Moreover, the search loop is parallelized matching each query spectrum on a separate thread.

After ranking all candidate spectra with the fragment index, we reevaluate the top hits, tracking all kinds of statistics (shared peak count, dot bias, an f-value equivalent as seen in SpectraST and b-y ion score, etc.). A refined *bias-adjusted similarity* measurement, which resolves m/z bins by modeling peak intensity spread with a Gaussian bell curve, determines the X highest scoring library spectra. X, the number of output PSMs per query spectrum (X*>*0), is a parameter defined by the user. The treatment of scoring functions for the case of simulated mass spectra and a step-by-step walk-through of the search loop including the implementation of SIMD with intrinsic C++ functions are discussed in the supplementary text.

Once all scheduled queries are performed, the resulting PSMs from all the partitions are concatenated and sorted by query ID. Matches assigned to the same experimental spectrum cluster together, and again only the top X ranked matches are retained, if a query was carried out in multiple partitions.

### 2.3 Data preparation

We evaluate the performance of Mistle on two common mock communities, 9MM (Tanca *et al*., 2013) and SIHUMIx (Krause *et al*., 2020). For the latter, we follow the recently published CAMPI study (Van Den Bossche *et al*., 2021), such that the evaluation is on par with the current metaproteomic benchmarking standard.

Protein sequence databases are re-utilized from Tanca *et al*. (2014) and the CAMPI study. Four original search files for 9MM and two large search files from CAMPI are selceted for the comparison. A summary of the two microbiomes and the data is found in the supplementary text.

#### 2.3.1 Spectral library construction

The protein sequence databases are digested using EncyclopeDIA (Searle *et al*., 2020) with a mass range of 400 to 1500 Da, charge states 2 to 4, and up to two missed cleavages. The normalized collision energy (NCE) is left at default value 33. Then, a locally installed version of Prosit, downloaded from https://github.com/kusterlab/prosit in 2019, is used to predict MS/MS spectra for all peptides and charge conformations in the peptide list. This way, the only modification considered is Cysteine Carbamidomethylation, which is fixed. A decoy library, when necessary, is created using DecoyPyrat (Wright and Choudhary, 2016) with minimum peptide length 7, and the downstream procedure is executed identically. Note that the SIHUMIx database already contains decoy sequences. Here, the database is split into two separate sets instead, before digestion and prediction. For 9MM, the contaminants database cRAP is appended, downloaded from the GPM FTP site http://ftp.thegpm.org/fasta/cRAP/ in December 2021. Additionally, the human proteome, downloaded from Uniprot Proteomes (Proteome ID: UP000005640), is added as an entrapment database to the target sequences. Again, we produce corresponding decoys with the method described above and digest and predict the spectral libraries accordingly. All of this results in 38 GB of spectral data (from Prosit in .msp format) for 9MM and 23.3 GB for SIHUMIx.

#### 2.3.2 Search set-up

The *mistle-build* indexing algorithm is applied to the data creating a searchable index with 64 search partitions in condensed binary format for the target and decoy library. The four experimental 9MM files are then searched using the *mistle-search* algorithm with 10 ppm precursor tolerance and 0.2 Da fragment tolerance (bin size), as suggested by Tanca *et al*. (2013). The two experimental files from the CAMPI study are searched in the SIHUMIx library with 10 ppm precursor tolerance and 0.02 Da fragment tolerance as was done by Van Den Bossche *et al*. (2021).

We conduct the exact same searches with SpectraST and msSLASH (Wang *et al*., 2020)on the target and decoy libraries, and with MSFragger given the original protein sequence databases. MSFragger 3.4 was used via the FragPipe pipeline. SpectraST version 5.0 was installed together with the TPP v6.0.0 software (Deutsch *et al*., 2015). Precursor and fragment tolerances are set as described above, peptide mass ranges are defined accordingly, and modifications are set to carbamidomethylated Cysteine only, to ensure a fair comparison. Aside from that, all tools run with default parameters. All pre-processing steps and mass calibrations are allowed. Since SpectraST and msSLASH accept precursor tolerance only in absolute values, we set it to 0.015, so that it considers all candidates for the largest peptides (10 ppm of 1500 Da). We were unable to produce interpretable results using msSLASH. Even with creating a peptide index and tracing back IDs, the output was not useful. Therefore, msSLASH is only considered when benchmarking run time and memory consumption.

#### 2.3.3 Quality control

False discovery rate (FDR) estimation using target decoy competition is put in place as primary technique to ensure quality of identification. For separate target and decoy searches (Mistle and SpectraST) the results are first merged, retrieving only the top scoring hit, either from the target or the decoy library. The FDR is then estimated from the number of target peptides *N*_target_ emitted at any scoring threshold *t* and corresponding the number of decoys *N*_decoy_, as a measure of false discoveries among them:

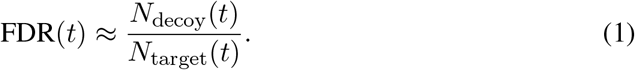

Afterwards, the FDR estimate is validated using *Homo sapiens* protein sequences as entrapment database. Target peptide identification that are unmistakably ascribed to human proteins are deemed false positives and an entrapment false discovery estimate can be computed by:

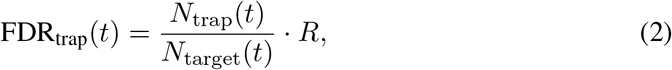

where *N*_trap_ is the number of entrapment peptides among the target identification and *N*_target_ are all target identifications at scoring threshold *t*. R is the ratio of database sizes, i.e. target database size over entrapment database size (number of peptides).

### 2.4 Run time and memory consumption evaluation

All searches are performed on a Debian 5.10.113-1 system with an Intel(R) Xeon(R) Gold 5120 CPU @ 2.20GHz and RAM of type DDR4, and with the data residing on an SSD. Eight threads were provided for each tool to make use of.

## 3 Results

### 3.1 Run time and memory performance

Time and hardware resources can become a critical factor when looking at large metaproteomes, covering thousands of species. For the moment, we evaluate run time and memory performances of all search software on the two small lab-assembled microbiomes, 9MM and SIHUMIx. Feasibility for larger databases is discussed later on.

#### 3.1.1 Run time

MSFragger is currently one of the most popular and time-wise best performing database search algorithms. We use MSFragger as representative of sequence database search algorithms in contrast to the spectral library search algorithms we evaluate against. Note that spectral library search faces inherently different bottlenecks, e.g. data loading and continuous index construction, since it does not have the whole database information immediately, unless it loads every spectrum into RAM. At the same time, no sequence processing is required, e.g. protein digestion. As for the spectral library search engines, we compare Mistle to SpectraST, as it is a stable and popular option among spectral search software. Additionally, we benchmark msSLASH, which has been recently developed and introduces massive run time improvements by using Locality-Sensitive Hashing (Wang *et al*., 2020).

Figure 3 compares the time required to search all experimental files between all four algorithms for both datasets. Mistle outperforms SpectraST by a factor greater than 10, and is about double as fast than msSLASH, for both spectral libraries. The gain in performance puts Mistle at a level comparable to the database search algorithm MSFragger, which is still a couple of times faster. Reasons for that are: i) the large index construction time, which is half of the total run time measured, since it requires multiple I/O operations to read MS/MS spectra, and repeatedly save their fragments to index partitions, and ii) a current lack of optimization for multiple search files that require reloading index partitions between runs. Combining consecutive searches into a single query immediately speeds up the process. We test this by concatenating the four 9MM search files into a single search file and analyze it with Mistle. While still querying the exact same experimental spectra, the search time reduces from 21 minutes down to 7 minutes, which is almost as fast as MSFragger. Also, note that the fragment index of Mistle needs to be constructed only once for each spectral library. Hence, when more files (or spectra) are searched, the run time reduces in relation to the other tools. Indexing time is indicated by the striped section in Figure 3 for Mistle and SpectraST. Note that for predicted spectral libraries, the indexing time gets overshadowed by the construction and prediction process of the spectral data itself, which is much more time consuming.

**Figure 3:**
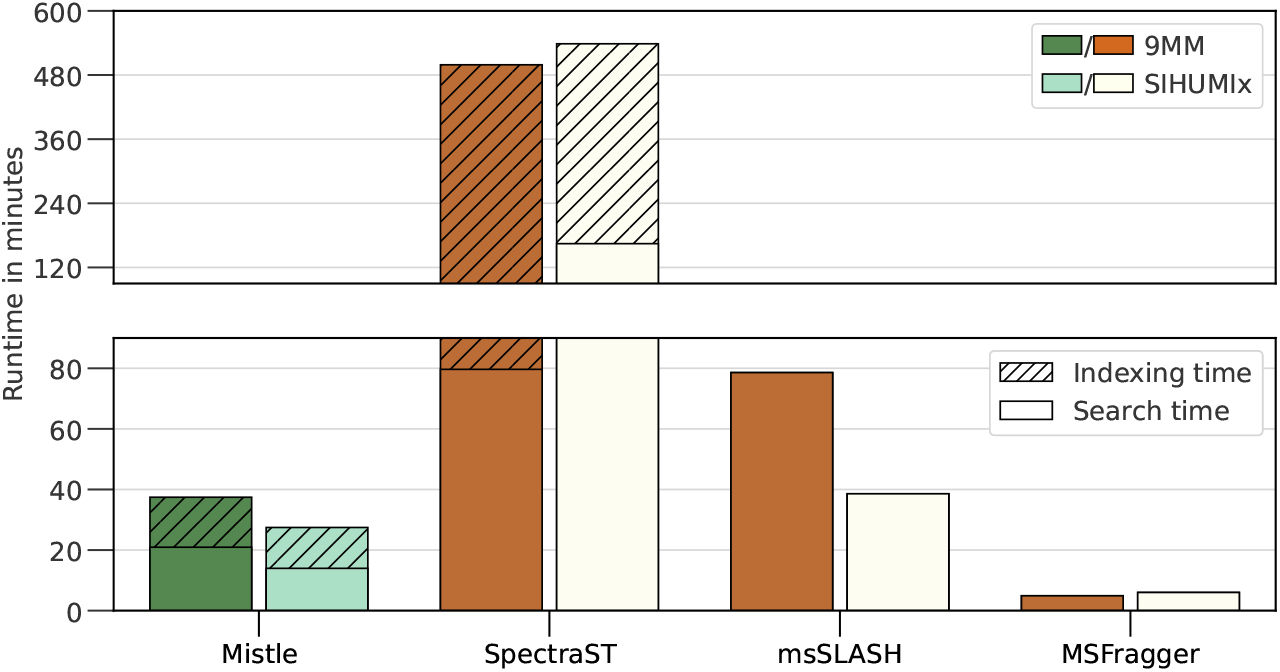
Run time measured for all 9MM runs and all SIHUMIx runs. Mistle is highlighted in green. Indexing time is indicated by stripes, whenever a separate index instance is constructed and saved to disk as intermediary step. Mistle performs the queries faster than any spectral library search engine. MSFragger is more than 4 times faster than Mistle. Half of the run time of Mistle goes into index construction.

Additionally, we investigated the time spent in distinct parts of the search loops, finding that *mistle-search* (lower bar in Figure 3) uses more than 90% of its time for loading the spectral index and constructing data structures. Conversely, Mistle spends ¡10% of the time performing spectral queries, i.e. candidate search in the precursor index, fragment matching, and the final rescoring of top-ranked candidates. This demonstrates a current bottleneck at index loading operations that could benefit from further optimizations in the future, such as the use of memory mapping.

Still, time-wise the fragment index search of Mistle introduces major improvements to spectral library search and brings it into a feasible reach compared to sequence database search.

#### 3.1.2 Memory

We analyze the memory requirements for all software, measured by the peak memory consumption across all runs for 9MM and SIHUMIx, respectively. In all cases, the target library was composed of the respective metaproteome, the human proteome and the contaminants database with the decoy library matching those. Figure 4 depicts the memory consumption in Gigabyte (GB). We find extraordinary memory improvements by the index partitioning and search scheduling method implemented in Mistle compared to all other search software, being around an order of magnitude more memory efficient. Compared to SpectraST, Mistle requires less than 18 times the amount of RAM, performing the exact same task. Mistle effectively constructs a fragment index and performs searches in a 38 GB large MS/MS library (9MM) with less than 3 GB RAM at all times, enabling queries on low performance computers, e.g. home laptops. We provide detailed information on how index partitioning and search scheduling affect run time and memory consumption in Figure 5. While the run time remains relatively constant, even seeing slight improvements with increasing partition count, the drop in memory consumption is eminent. The complete index (one partition) is comparable in size to MSFragger’s fragment index, though it holds additional information, such as peak intensity values. However, with the use of more than one partition, the RAM usage decreases according to the partition size. The usage converges to roughly 2 GB when using hundreds of partitions. This is likely the cached size of precursor index and query spectra together, while the fragment index size gets arbitrarily small by the partitioning. We use Mistle with 64 partitions in this study as it significantly reduces memory consumption with a stable run time, but the theoretical optimum is not reached with 64 partitions for this dataset. We test imposing similar memory constraints on MSFragger, by using its split database function with 64 chunks and maximum 3GB

**Figure 4:**
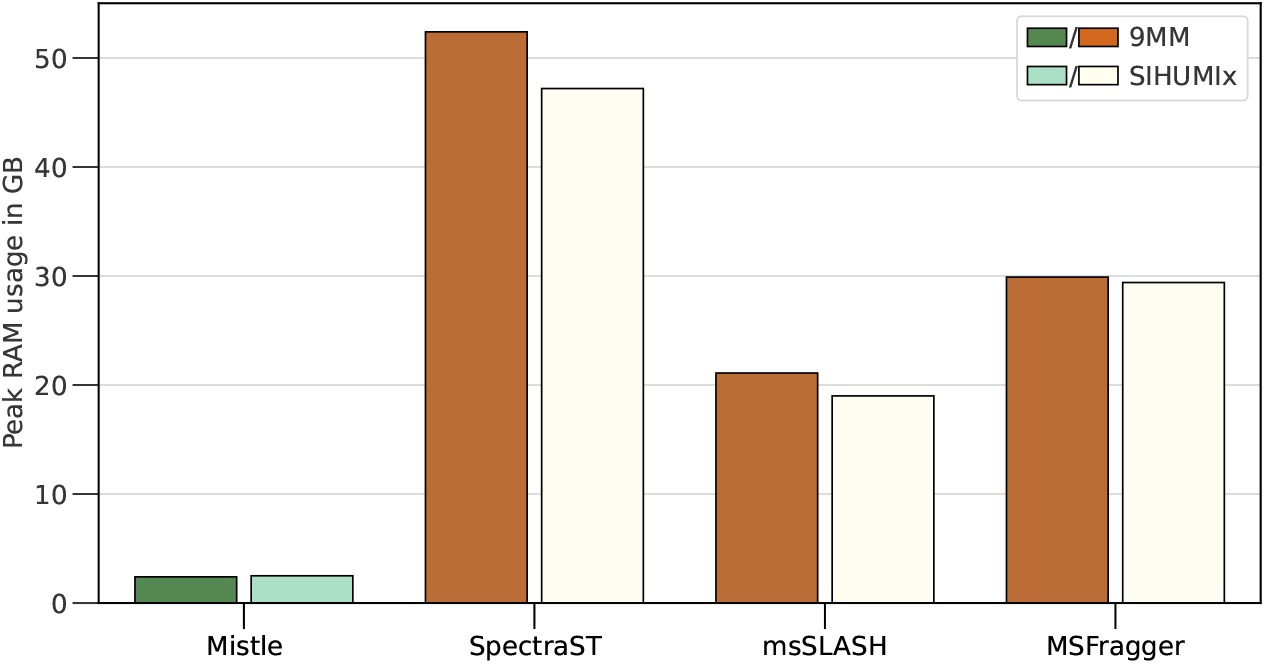
Memory consumption measured over all 9MM runs and all SIHUMIx runs. Mistle performs much more RAM efficiently than all other tools, with almost a 20-fold decrease in memory usage when compared to SpectraST and up to 10-fold decrease when compared to msSLASH, and *>*10-fold decrease when compared to MSFragger.

**Figure 5:**
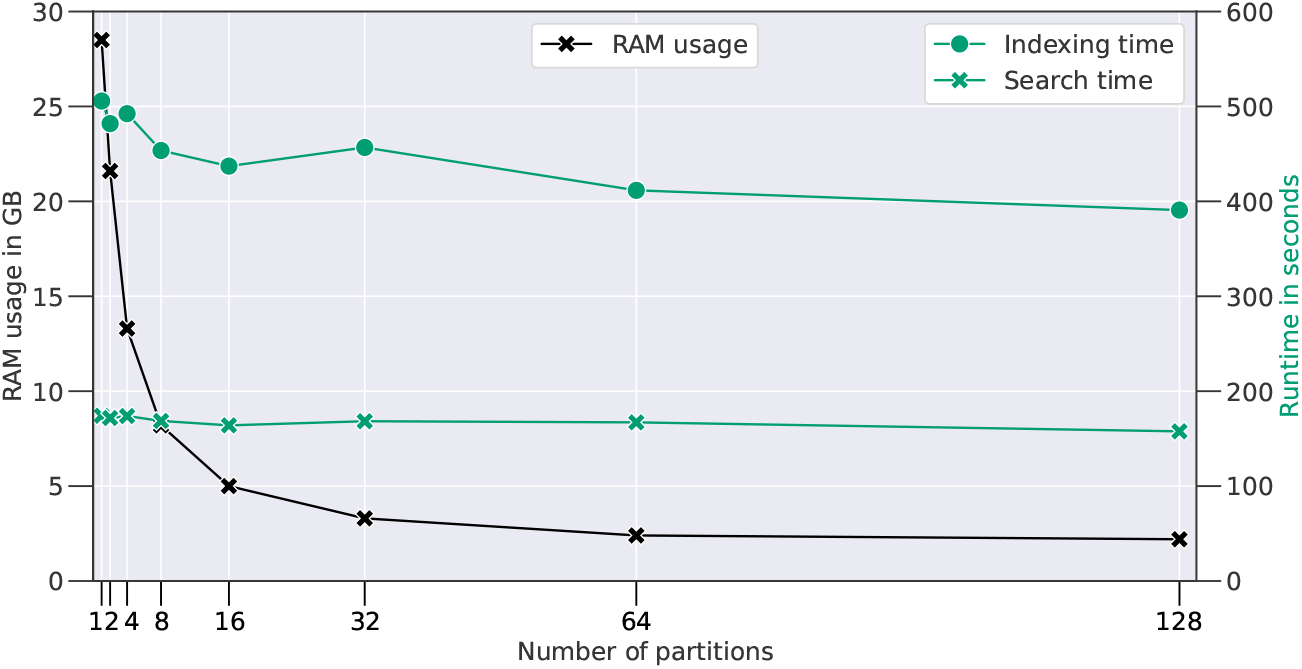
Memory consumption (black) and run time (green) shown for increasing numbers of partitions of Mistle’s fragment index for the 9MM target library. Peak RAM was measured, and the run time is divided into the time required to build the fragment index (dots) and search time (crosses) for the largest search file (9MM Run 1.mgf).

RAM allowed. Aside from the program frequently stopping with non-zero exit code, the database search takes up to 8 times longer than before, when running through. This again emphasizes the effectiveness of Mistle’s partitioning algorithm, advertising for a similar implementation in MSFragger, if possible.

Figure 5 also highlights the imbalance between search time and index construction time, as mentioned before. For a single search file, index construction takes almost three times longer than the spectral matching and ranking process, rendering Misle less efficient when the library is only queried once. On the flip side, many or particularly expansive MS/MS runs evaluated against the same metaproteome database benefit from an excellent search time on individual runs, where the fragment index (and spectral library) only needs to be constructed at the outset of an analysis.

### 3.2 Peptide identification

We investigate the peptide identification rate for PSMs and unique peptides, of the predicted spectral library approach using Mistle and SpectraST, and compare it to database search via MSFragger. Only the discriminant score from each search engine is taken into account and the false discovery rate (FDR) is estimated with the target decoy approach, according to equation 1. The total number of significant PSMs is plotted over the corresponding FDR estimates for only the first search file of each database in Fig. 6 (top left). The other search files show a similar picture. In terms of number of identifications, Mistle yields the most matches for 9MM (Fig. 6a) with 11,845 PSMs at the critical 1% FDR level (vertical dotted line). Here, MSFragger finds 7,921 at the same FDR threshold. This difference is also reflected in the unique peptides identified throughout all searches, as seen in the upsetplot in Fig. 6a. The intersection of identified peptides are displayed for each tool with every other. Mistle, finds the most unique peptide. Noteworthy though, the spectral search approach, i.e. Mistle and SpectraST, yields a large distinct overlap of *>*2000 peptides, suggesting a non-negligible number of unique peptides only significantly identifiable when taking peak intensity predictions into account.

**Figure 6:**
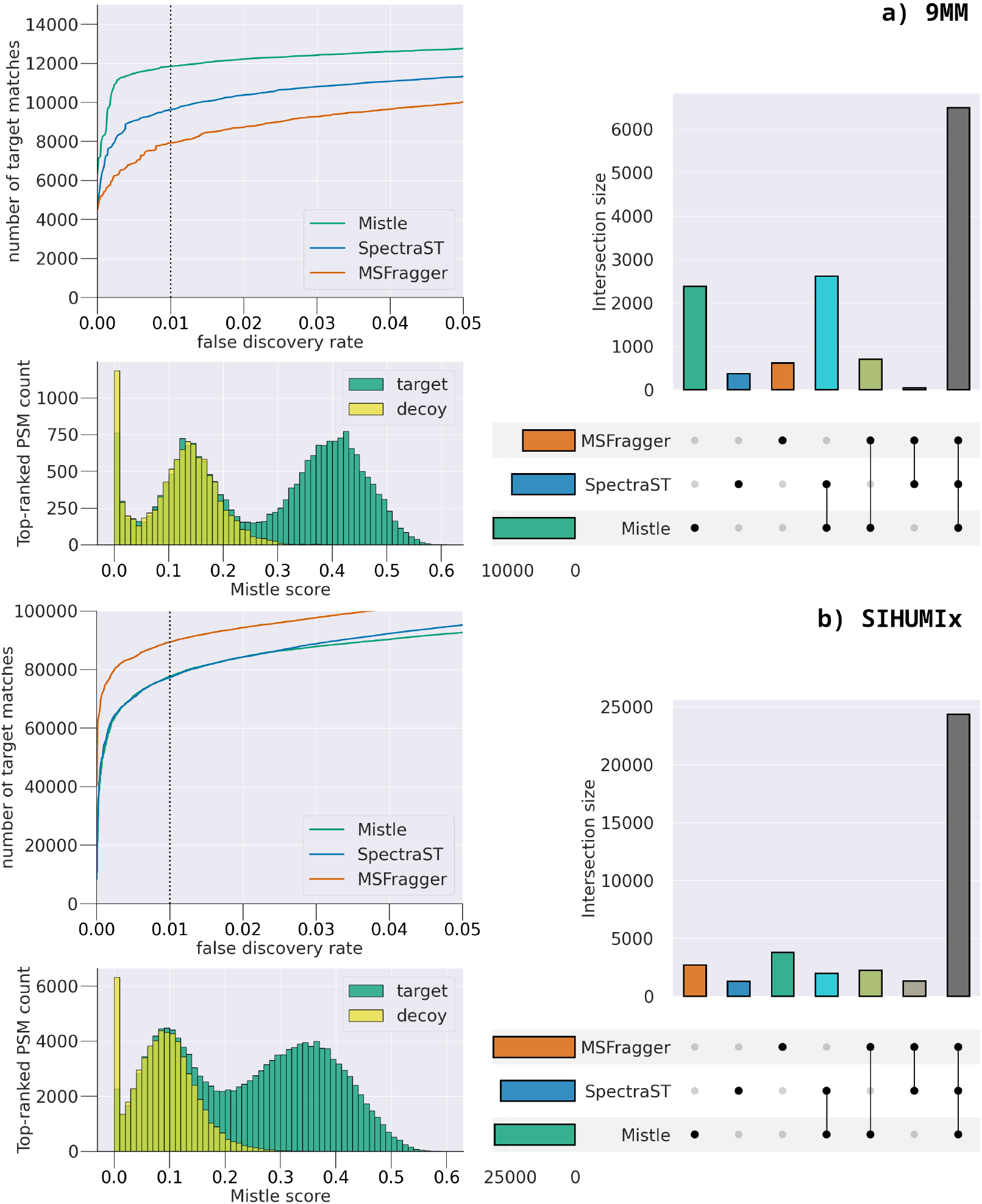
Statistics on peptide identification for 9MM (top panel) and SIHUMIx (bottom panel). The left side shows the number of PSMs identified over the estimated FDR for all tools, exemplary for first search file (9MM FASP.mgf and S05.mgf, respectively). Below, Mistle’s target (green) and decoy scores (yellow) for top ranked PSMs are plotted for the same query. A bimodal distribution for target scores can be seen for both databases. On the right the total set of uniquely identified peptides (all search files) are shown as an upset plot for all kinds of intersections; corresponding Venn diagrams can be found in the supplement.

For the SIHUMIx database (Figure 6b) the roles are reversed, with MSFragger finding a large number of PSMs (89,403 at 1% FDR), roughly 10,000 more than Mistle and SpectraST. Half of these additional PSMs can be explained by the mass calibration and optimization steps performed by MSFragger, which yields an additional 5,200 PSMs (6% of the total), compared to when turning off this feature. Mistle and SpectraST do not offer such a specific feature. In terms of unique peptides, we see a large overlap between all three tools. This makes a lot of sense, considering that *>*400,000 MS/MS scans have been queried. The distinct intersection between Mistle and SpectraST is less pronounced here, but this may also arise from both spectral search engines being less sensitive for the SI-HUMIx runs. Even though the total number of output PSMs is reduced for Mistle, almost the same number of unique peptides are identified compared to MSFragger, potentially indicating these peptides to be truly present. Compared to the CAMPI study, we observe fewer significant PSMs at 1% FDR: ¡100,000 for all search engines and files, whereas ca. 120,000 were identified in the CAMPI study. However, our library set-up is more restrictive in terms of modification and peptide size - compare section 2.3 with Van Den Bossche *et al*. (2021) - making it hard to judge the difference in raw numbers. Additionally, the statistics for S05 and S06 search files against the SIHUMIx reference database in the CAMPI study were attained by database search with X!Tandem (Craig and Beavis, 2004), which uses a refinement step, likely yielding additional hits.

For both the metaproteomes queried, the scores Mistle assigns to matched target peptides follow a bimodal distribution, with the decoy scores following one of the modes (bottom left histogram in Figure 6a and b). This is an indication of a clean separation of true and false target PSMs by the bias-adjusted scoring function used in Mistle. If such behavior manifests throughout more and specifically larger datasets, omitting target decoy competition altogether might be a reasonable option. A mixture model approach as for example discussed by Nesvizhskii (2010) comes to mind. Dropping decoys sequences would reduce resources needed for library construction, fragment indexing and search, coming in very handy for large-scale metaproteomic studies.

### 3.3 Quality Control Validation

Peak intensities predicted by Prosit provide a detailed picture of which candidates are best suited to explain an experimental spectrum. To investigate how this improves the quality of identifications, we set up an entrapment approach to verify FDR and present examples showcasing where our approach produces a better match.

Figure 7a shows such an example MS/MS scan from the SIHUMIx query (top of the mirror plot) paired with the spectral prediction from Prosit (bottom) of its identified peptide by Mistle, and Figure 7b shows the same spectrum matched by MSFragger. The near perfect overlap between b and y ion intensities suggests that Mistle identified the correct peptide, while the identification by MSFragger on the same spectrum is an apparent mismatch to a decoy peptide. This is only obvious given the spectral predictions by Prosit, as the theoretical b and y ions cover only a fraction of all peaks in both cases, amounting to similar peak intensities. Note that using Prosit-Rescore to post-process MSFragger’s search results might eliminate such false discoveries, but still forgo the correct PSMs, unless many lower ranked matches are rescored as well. Of course, this would dampen the run time significantly.

**Figure 7:**
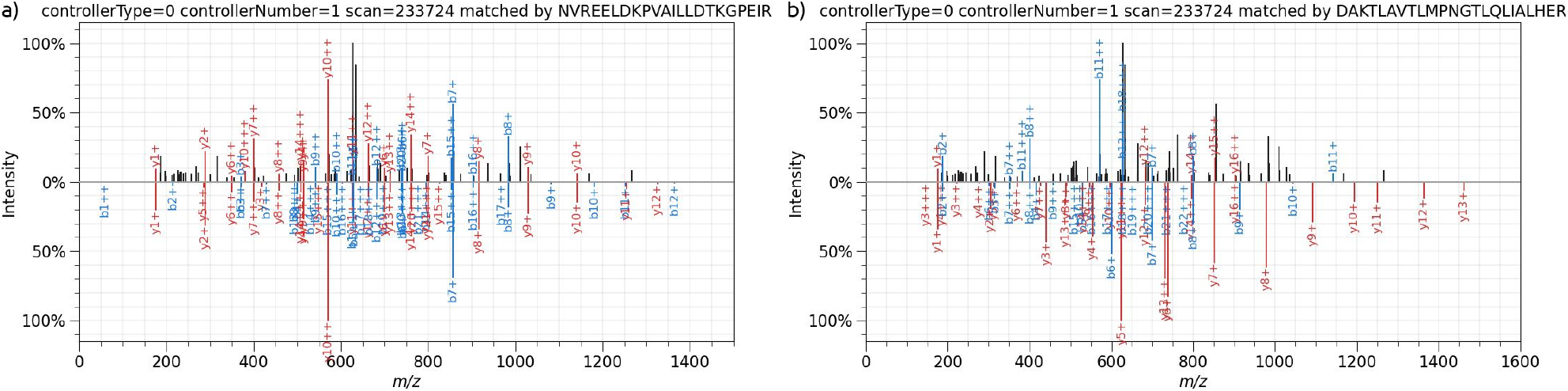
Mirror plot of a PSM identified by Mistle a) and MSFragger b). In each case the top spectrum is the same experimental spectrum, scan 233724 of S05.mgf file and the bottom spectrum is the matched peptide spectrum from the SIHUMIx database with the peak intensities predicted by Prosit. Mistle finds a reasonable candidate. MSFragger identifies a decoy sequence.

While FDR estimation via target decoy competition is an established method to ensure quality of peptide identifications, we like to assess the level of confidence we can put into the estimate. Feng *et al*. (2017) uses a human entrapment database as a standard, to compare quality control methods against, for sample sequences from *Pyrococcus furiosus*. Similarily, we set up our target sequences to contain both, sample sequences (9MM or SI-HUMIx) and *Homo sapiens* sequences. The corresponding entrapment FDR is computed using Equation 2.

Figure 8 shows the entrapment FDR over the target decoy FDR estimate, as a range (minimum, maximum and average) across all 9MM runs. Along various thresholds, Mistle estimates the FDR well using the target decoy approach compared to the entrapment FDR, especially at the critical 1% FDR threshold, with small variation between runs. The more lenient the FDR control is chosen (*>*1%), the more Mistle seems to overestimate the target decoy FDR slightly. However, this only exposes the scoring threshold as too stringent, yielding fewer PSMs, but still guaranteeing a correct or even smaller FDR. Comparable results are seen for SpectraST. Overall, this confirms that the spectral library approach produces PSMs with a stable FDR estimation, indicating the results are reliable. MSFragger, in contrast, slightly underestimates the FDR with the target decoy approach when looking at the corresponding entrapment FDR estimate, across the full range (0 to 5% FDR). This is discerning, since the output may contain more errors than specified, rendering the quality control measurements to be questionable. Also, more variation across the runs can be seen for MSFragger, as the increased differences between maximum and minimum entrapment FDR illustrate.

**Figure 8:**
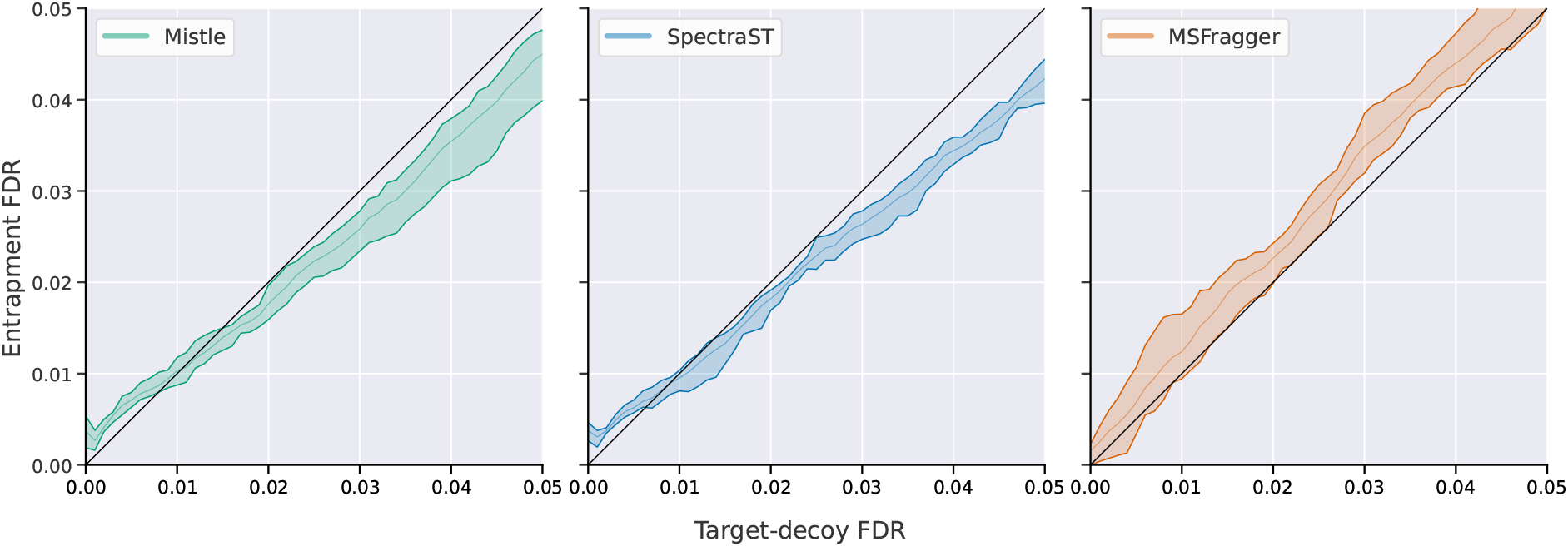
Entrapment FDR over target decoy FDR measured across for all four 9MM runs. The range is displayed by the colored area given by the maximum and minimum entrapment FDR for any target decoy FDR interval with average being the inner line. Mistle is green, SpectraST blue, and MSFragger orange. The black slope-1 line indicates the desired scenario, where the two FDR estimates align perfectly.

Searching the SIHUMIx database, all tools produce PSMs with robust FDR estimates, which are slightly too conservative, especially in the case of SpectraST. A corresponding figure can be found in the supplement.

## 4 Discussion

Coping with complete predicted libraries for metaproteomics, covering more than 10 million peptide MS/MS spectra, proves to be a challenging task for spectral search software. Especially, analyzing the large candidate space in RAM is very demanding and might just fail looking at more diverse microbiomes. We provide proof of concept that our approach works well for two mock-communities, turning all peptides from their sequence database into MS/MS spectra. Despite the large data sets involved, the presented search algorithm, Mistle, is extremely memory efficient due to an effective index partitioning technique. The memory requirements of Mistle are an order of magnitude smaller than those of all other software. In terms of run time, Mistle is faster than other spectral library search software, and stays close to the ultra-fast database search algorithm MSFragger. Even though Mistle cannot quite match the run time of MSFragger, the challenges faced by each approach are quite different, and the improvements introduced by our algorithm are nonetheless significant.

Investigating peptide identification, we find cohesive performance between Mistle and SpectraST identifying adequate numbers of PSMs (see Figure 6). The top ranked hits emit the desired bimodal distribution of target scores. An elevated peptide overlap between SpectraST and Mistle reinforces the idea of spectral matching being able to identify peptides that standard database search cannot. Delving into this, we presented an example spectrum visibly attaining a much more reasonable match using our approach, when compared to MSFragger, which identifies a decoy peptide (see Figure 7). Entrapment sequences added to target proteins to measure FDR estimate stringency reveal enhanced precision when identifying peptides with the spectral matching approach (see Figure 8). As we evaluate the direct performance of the different search approaches, post-processing tools like Percolator (Käll *et al*., 2007) are omitted completely. However, it stands to reason that Percolator can make good use of the myriad of scores and statistics that are being tracked by Mistle. This may improve identification rate and accuracy even further. In conclusion, we prove the applicability to common lab-assembled studies and have reason to believe the workflow will perform well for even larger metaproteomics studies, if set up properly.

Currently, the main shortcoming resides in building the spectral library as an in-between step, which is resource intensive (time, and disk space). Additionally, loading times from disk take up more than 90% of the total search time and are the reason for slightly increased run time compared to state-of-the-art database search methods like MSFragger, regardless of the optimizations we put in place. A way to mitigate long loading times is to distribute search tasks among several servers, each permanently keeping an index partition in RAM. As a beneficial side effect partitions can then be queried in parallel without any I/O operations. Of course, this is resource demanding in its own way.

While the workflow produces satisfactory results no matter which spectral search engine is used, Mistle has an excellent trade-off between run time and memory consumption and outperforms SpectraST in that aspect by far. Mistle is best used for repeated scans on the same metaproteomic environment, like for instance SIHUMIx, such that the time-consuming spectral library and search index are constructed only once. The sequence database and parameters can be chosen generously to be very comprehensive, and the performant search time comes to shine on multiple MS/MS runs against the same library. The low memory consumption even makes Mistle feasible for studies on low performance machines, e.g. home laptops, but also allows much larger protein databases to be analyzed, where the competing tools reach their limit in feasibility. Of course, Mistle, being a spectral search engine at heart, can be used on any experimental spectral library, too. It is especially suited for large spectral datasets and provides various options and statistics to the user.

There are small qualitative differences observed between SpectraST and Mistle. They arise from different pre-processing steps and the scoring functions used, e.g. neighboring bin matching for SpectraST and peak matching using a gaussian distribution by Mistle. Turning off most pre-processing features and using the native dot product for scoring produces nearly identical results between SpectraST and Mistle. At this point, we conclude that the differences in PSM scoring play a minor role, when looking at the overall identification of unique peptides in the samples. Figure 6 demonstrates a large overlap in the findings of both spectral library search engines (*>*70% of unique peptides are shared for the two studies).

With our tool, we open the door to investigate much larger metaproteomes, e.g. the human gut microbiome, with the help of predicted spectral libraries. In an ideal setting, spectral library prediction is set up to cover the entire metaproteome comprehensively with carefully selected parameters in accordance with the wet lab. Then, the effect of treatments, different samples or patient groups can be perpetually analyzed by spectral search with Mistle producing reliable peptide identification in the large search space at fast rate, and without being memory intensive. Higher level analysis can build upon these identifications. Nonetheless, the construction of predicted libraries remains time and resource consuming, easily going into terabytes of spectral data for the human gut. We are curious to see this being put into place in future studies.

## Acknowledgements

We thank our colleagues Tanja Holstein and Sasan Amariamir for stimulating discussions and insights that assisted this research.

## Supplementary Material

### 1 Summary

This is the supplementary material to the article *Mistle: bringing spectral library predictions to metaproteomics with an efficient search index*. Additional explanation of the methods and datasets are provided. Moreover, supplementary figures supporting the study are attached in Section 3.

### 2 Methods

In this section we provide additional information to some of method deployed in the main article, and elaborate the data processing steps. This serves as ancillary information, and is only fully comprehensible in combination with the main text.

#### 2.1 Continuous index construction

As explained in the main article, the fragment index needs to be constructed continuously during the reading process of the spectral library.

This goes as follows: Each library spectrum is read one after the other, precursor information is added to the precursor index and all their peaks are streamed as fragment triplets to the corresponding index partition in binary format on disk. This is parallelized by having a single thread dedicated to reading while the rest are occupied by formatting and writing tasks. At this point, fragments are unsorted and not binned, but assigned to their partition. After the library has been read entirely, the precursor index is sorted by precursor charge and m/z, and the ID-to-rank mapping is established by a linear scan over the precursor list. Then, every partition is loaded again, one at a time, and fragments are sorted based on parent ranks, accessible via the ID. Afterwards, the partition is saved to disk in binary format. Binning only happens during the search, based on the user-specific fragment tolerance.

#### 2.2 Search Loop

Let *Q* be a query spectrum with precursor m/z *mz*_*Q*_ and a set of peaks *P*_*Q*_. We identify the best PSMs by computing the dot product to the reference library spectra, carrying out the following steps:

1. First, the range of library candidate spectra that lie within the precursor mass tolerance *t* is computed using the precursor index. A binary search on the sorted precursor entries swiftly finds the lower bound *L* and upper bound *U*, which is the rank of the first and last library spectrum with a precursor m/z ∈ [*mz*_*Q*_ − *t, mz*_*Q*_ + *t*].
2. Next, a scoring vector (scores) of length *U* − *L* is allocated and initialized with all zeros. This way, the score of each candidate spectrum can be accessed from its rank *R* by subtracting *L*.
3. Every peak *p* = (*mz*_*p*_, *I*_*p*_) ∈ *P*_*Q*_ is searched in the fragment index by first determining the fragment bin matching its ion mass *mz*_*p*_. Recall that inside each bin the fragments are stored in order of their parent ranks. A binary search then identifies the first fragment *f* = (*mz*_*f*_, *I*_*f*_, ID_parent_(f)) with a parent rank *R* greater than or equal to *L* (rank of the first candidate). *R* is derived from ID_parent_(f) via the ID-to-rank mapping.
4. The fragment intensity *I*_*f*_ is multiplied with the query peak intensity *I*_*p*_ and the product is added to the score of the fragment’s parent:

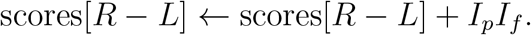

The process is repeated with the next fragment in the bin until a fragment with parent rank greater than *U* is reached. This marks the end of candidates for that fragment bin.
5. At the end, the scores equal the dot products between search spectrum *Q* and every candidate library spectrum covered by the partition. We rescore the best-scoring library spectra with an elaborate scoring function (see section 2.2). Then, the X highest scoring library spectra are selected, and the corresponding peptides are returned as PSMs to *Q*. X, the number of output PSMs per query spectrum (X*>*0), is a parameter defined by the user.

The search function is parallelized matching each query spectrum on a separate thread. After all scheduled queries are performed, the resulting PSMs from all the partitions are concatenated and sorted by query ID. Matches assigned to the same experimental spectrum cluster together, and again only the top X ranked matches are retained, if multiple partitions produced hits for the same query.

##### SIMD intrinsics

Single instruction multiple data (SIMD) is a type of parallel computation, which simultaneously performs an operation on packed groups of data, instead of on every single data point individually (Amiri and Shahbahrami, 2020). Since their introduction to general-purpose processors, first by Intel’s MultiMedia eXtensions (MMX) in 1996, it has been widely established as significant means to speed up performance (Amiri and Shahbahrami, 2020; Hassaballah *et al*., 2008; Zhou and Ross, 2002).

The search algorithm requires many successive multiplication and addition operations when updating parent scores with peak intensity products. Consequently, SIMD extensions are an eligible option to improve our run time. We use the Advanced Vector Extensions AVX2 and AVX512 architectures, which support the fused multiply-add arithmetic operation (for 256-bits in C++: *mm256 fmadd ps*) for floating-point vectors, thereby updating multiple parent scores in parallel. A schematic version of the workflow for a 256-bit register is depicted in Supplementary Figure 2.

Explicitly, this is done by broadcasting a single 32-bit float, the query peak intensity value, into the 256 or 512-bit register (for 256-bits in C++: *mm256 set1 ps*) and loading respectively 8 or 16 fragment intensities from the fragment-ion bin (using *mm256 loadu ps*). Then, the corresponding score values are inserted into another register. Finally, the multiply-add operation (using *mm256 fmadd ps*) is performed vertically on all 8 or 16 values at the same time: multiplying the query intensity with the individual fragment intensities and adding the result to the corresponding scores, which are extracted afterwards. The AVX512 instruction set provides a *gather* instruction to quickly access the values from the scoring vector and a *scatter* instruction to put them back in place after the computation. Note that the fragment bins need to be adjusted in their format to enable swift loading of intensity values.

##### Spectral similarity scoring

SpectraST features an option to spread a fraction of the peak intensity to neighboring bins to account for slightly shifted peaks (Lam *et al*., 2007). The inherent structure of the fragment index impedes matching such shifted peaks efficiently. To circumvent this problem, we resolve the bins after the initial search is performed. Top ranked hits based on the spectral dot product (Eq. 2 by Lam *et al*. (2007)) are rescored by another formula, which models peak intensity spread with a Gaussian bell curve put over the peaks and penalizes mass shifts as intensity decay along the curve. We define a similarity between a query spectrum *Q* and reference spectrum *R*, analogous to the dot product of binned intensities:

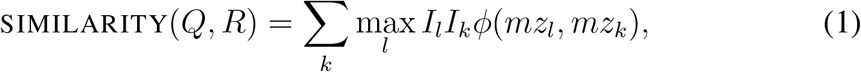

and

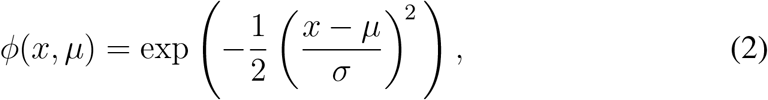

where *k* iterates over all reference peaks (*mz*_*k*_, *I*_*k*_) ∈ *R* and *l* over all query peaks (*mz*_*l*_, *I*_*l*_) ∈ *Q. ϕ*(*x, µ*) factors in the distance of *mz*-values given the standard deviation *σ* between peaks. We set *σ* to half the bin size. An analogous formula to the dot bias as provided by SpectraST (Lam *et al*., 2007) can be established:

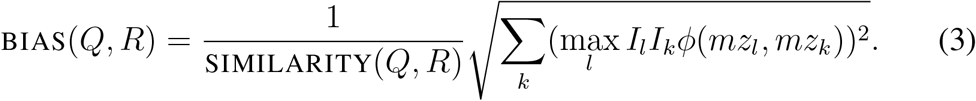

Mistle tracks numerous scores, such as the ion count, and the f-value used in SpectraST. For the purpose of this study, we simply deploy a bias-adjusted similarity measurement as: SIMILARITY (1 BIAS).

Note that as Prosit only predicts b and y-ion intensities, it might be ill-advised to score similarity based on all peaks. A correct match might achieve an inferior score, when additional ion-type peaks are present, which are not matched by the prediction. Therefore, we introduce a second similarity, which only considers similarity of matched peaks, which we call *b-y ion score*. Bias and bias-adjusted similarity are formulated equivalently on b and y-ions only. The final scoring for the evaluation of target decoy competition is the average between bias-adjusted similarity and the b-y ion version of the same formula. Mind that the *b-y ion score* alone might not provide a perfect distinction either, since it can disregard large portions of the experimental MS/MS spectrum. A false peptide might achieve a high score by matching well to small peaks annotated as b and y ions yet leaving most of the peak intensity unaccounted for. Thus, we opt for the average out of both spectral similarity measurements.

#### 2.3 Datasets

##### 2.3.1 9MM

In 2013, Tanca *et al*. investigated the effect of sequence databases used to query shotgun proteomic results for diverse microbial communities. They evaluate their findings on a lab-assembled mock community of nine bacterial and eukaryotic species: *Escherichia coli, Pasteurella multocida, Brevibacillus laterosporus, Lactobacillus acidophilus, Lactobacillus casei, Enterococcus faecalis, Pediococcus pentosaceus, Rhodotorula glutinis*, and *Saccharomyces cerevisiae*. The 9MM dataset has since been used to evaluate metaproteomic pipelines. We utilize the sequence database (9MM DB.fasta) and 4 search files (9MM FASP.raw, 9MM PPID.raw, 9MM Run 1.raw, 9MM Run 2.raw) provided in the original and the follow-up study, which can be found at http://www.peptideatlas.org/PASS/PASS00194 and http://www.peptideatlas.org/PASS/PASS00355 (Tanca *et al*., 2013, 2014). Raw files were convered using ms-convert from the ProteoWizard software (Chambers *et al*., 2012) with peak picking retaining the top 150 most intense peaks.

##### 2.3.2 SIHUMIx

The extended simplified human microbiota (SIHUMIx), established by Krause *et al*. (2020), is a model community of eight species from the human intestine that account for most of the typical metabolic activities in the human gut. The model allows consistent and reproducible in-vitro cultativation, making it ideal to investigate the effect of treatments to the microbiome (Schäpe *et al*., 2020). SIHUMIx consists of the species: *Anaerostipes caccae, Bifidobacterium longum, Bacteroides thetaiotaomicron, Blautia producta, Clostridium butyricum, Clostridium ramosum, Escherichia coli* and *Lactobacillus plantarum*.

As reference sequence database we use the one provided by the CAMPI challenge (Van Den Bossche *et al*., 2021), which already includes decoy and contaminant sequences (Pride Project ID: PXD023217). Searches are performed on two large search files (S05, S06) from the CAMPI study, which yielded the most identified PSMs without fractionation.

### 3 Supplementary Figures

**Figure 1:**
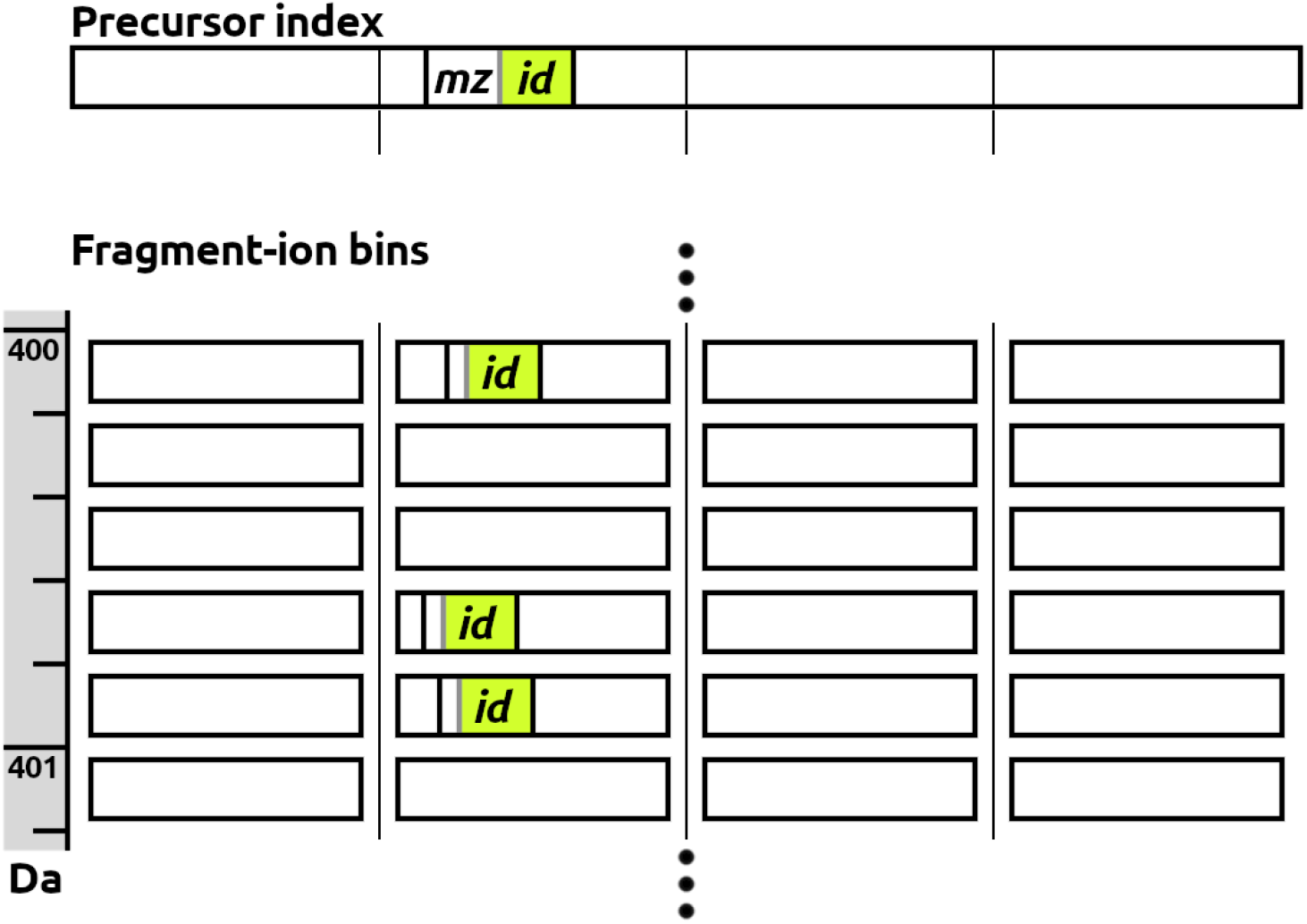
Schematic depiction of the fragment index partitioning into 4 subindices based on precursor index ranking. All peaks from a parent entry (highlighted on top) are listed inside the fragment sub-index (partition) determined by the parent’s m/z. For search and construction, only a single partition and the precursor index need to be loaded into RAM.

**Figure 2:**
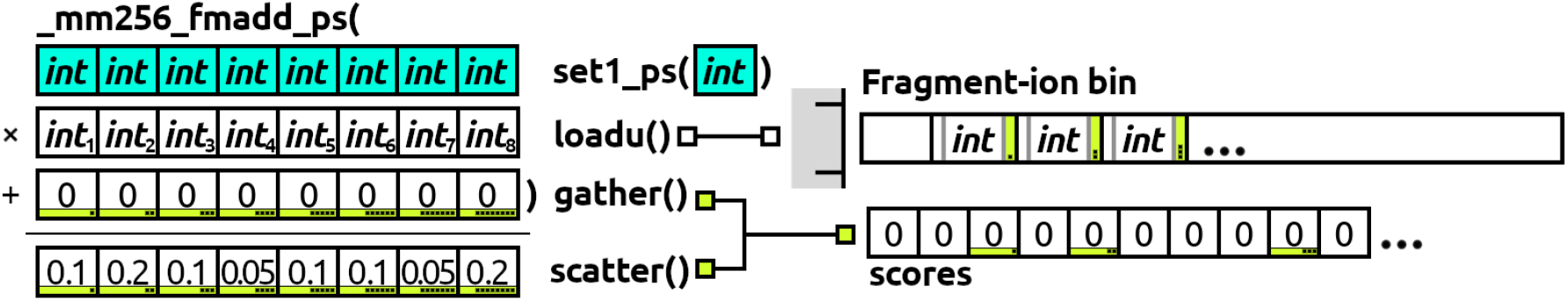
Visualization of AVX2 fused multiply-add operation performed vertically on 8 intensity values (32-bit floats). The process associated with data loading into and retrieving from 256-bit destinations is displayed for a single fragment bin. *Gather* and *scatter* instructions are only available for AVX512 compatible CPUs. This computation replaces step 4 of the inner search loop described in 2.2.

**Figure 3:**
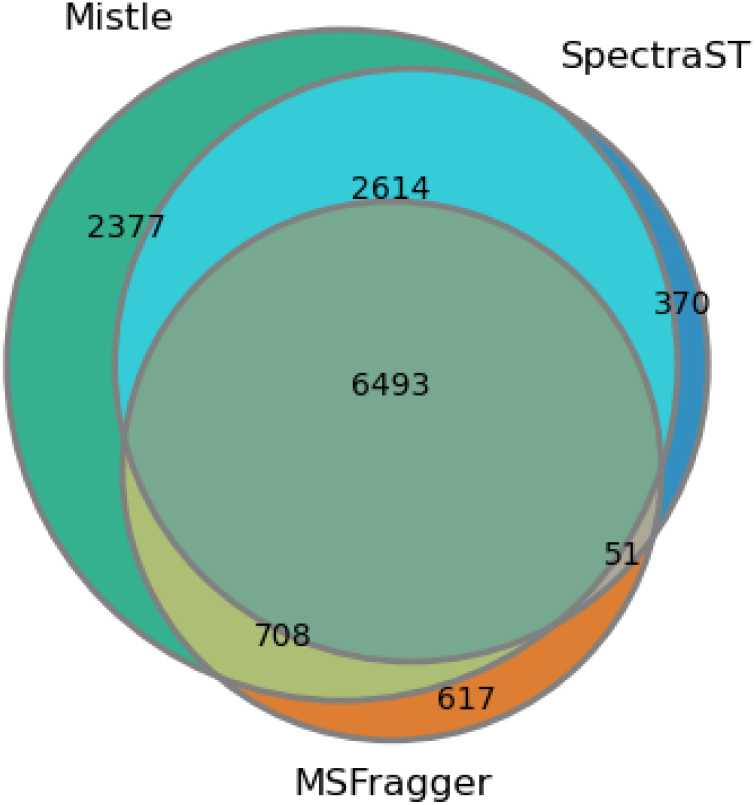
Venn diagram for 9MM dataset, corresponding to upset plot from Figure 6 in the main article.

**Figure 4:**
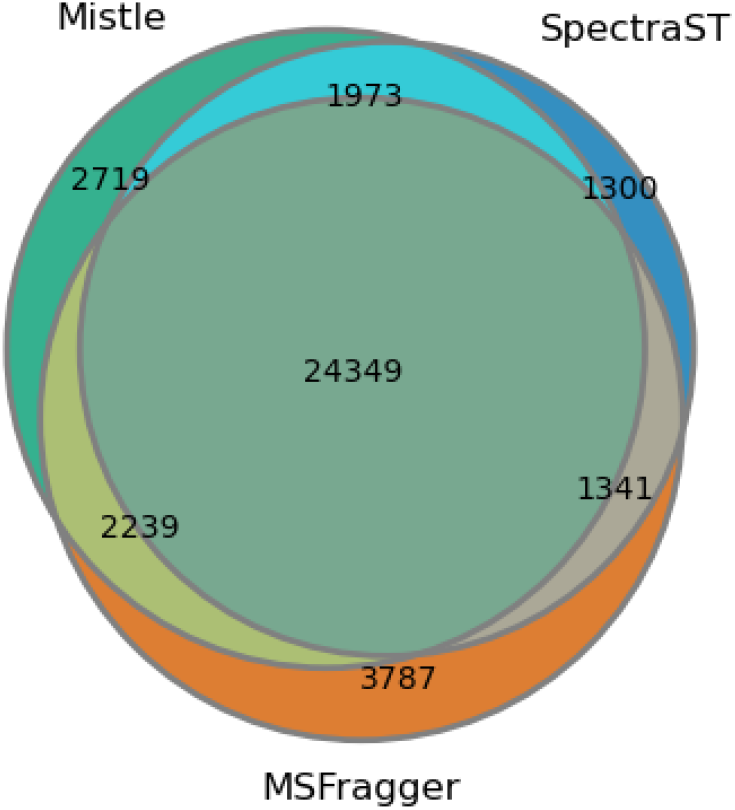
Venn diagram for SIHUMIx dataset, corresponding to upset plot from Figure 6 in the main article.

**Figure 5:**
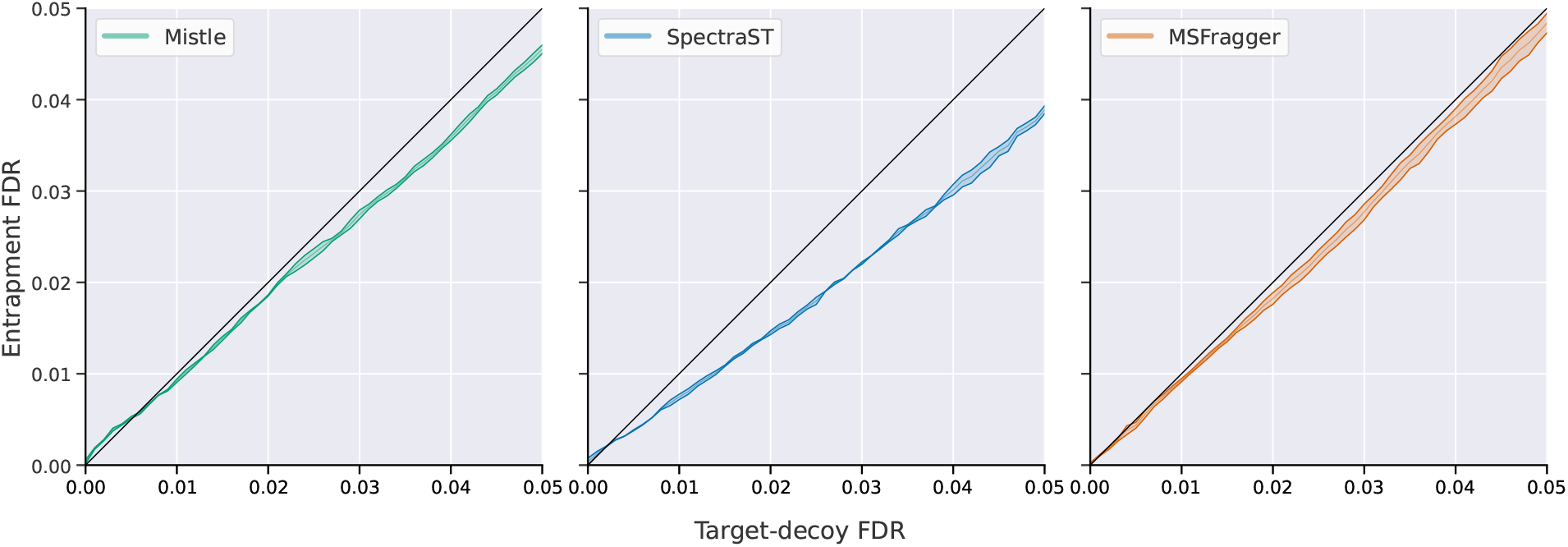
Entrapment FDR over target decoy FDR measured across the two SI-HUMIx runs. The range is displayed by the colored area given by the maximum and minimum entrapment FDR for any target decoy FDR interval with average being the inner line. Mistle is green, SpectraST blue, and MSFragger orange. The black slope-1 line indicates the desired scenario, where the two FDR estimates align perfectly.

